# Influence of the metallized coils on human leg blood circulation

**DOI:** 10.1101/2021.06.04.447146

**Authors:** Jurijs Dehtjars, Ksenija Jašina, Viesturs Larins, Aleksandrs Okss, Konstantins Pudovskis, Nelli Tolmača, Vijay Vyas Vadhiraj

## Abstract

The aim of the study was to find out if magnetic field generated by the human body affects a human blood flow. The idea is based on Lenz’s law where the blood flow induces an opposing alternating magnetic field (OAMF). In the experiment the OAMF will be modulated by repeating heart contractions (pulses). In an experiment with metallized coils it was found that wearing metal coils affects blood flow and it differs from when coils were not worn.

## Introduction

Chronic venous insufficiency (CVI) occurs when the venous wall and/or valves in the veins of the extremities are not working effectively, causing difficulties for the return of blood to the heart from the extremities [1]. The first disorders to occur are usually arrhythmia (the rhythm could become uneven or quickens) followed by the outflow of blood in the veins, especially the legs, which causes thrombophlebitis and the formation of varicose veins. Inadequate blood circulation leads to coronary heart disease, myocardial infarction, angina pectoris and other pathologies. Risk factors for the development of chronic venous insufficiency are believed to be age, genetics, gender, obesity, pregnancy, hormonal disorders, prolonged monotonous standing.

CVI is often overlooked by healthcare providers because of an underappreciation of the magnitude and impact of the problem, as well as incomplete recognition of its various manifestations of primary and secondary venous disorders [2]. However, based on a study conducted by Rabe et al., as of 2012 the worldwide prevalence of CVI in all its stages was 83.6% [3].

Therefore, an enhancement of blood circulation in the veins of legs to slow down the onset of CVI or reduce its symptoms [4].

Methods that employ magnetic fields to enhance blood circulation in humans have been used for centuries [5] exposed to low frequency (1-120 Hz) low induction (10^−7^ – 10^−5^ T) of alternating magnetic fields that increase in vascular area and shape [6], improve blood circulation [7][8] and one study performed on human subjects showed that the application of even a small low frequency magnetic field of about 1 mkT over a period of several minutes can lower blood pressure in human subjects [9]. Therefore, it is reasonable to assume that exposure to low frequency low induction alternating magnetic fields could have a therapeutic.

The most commonly used technique in magneto physiotherapy is so called pulse magnetic therapy [10]. For this type of therapy special devices are required. There is also magnetotherapy that provides on average 100 times less (low-frequency field: from 0-100 Hz) electromagnetic field in the tissues than in electrotherapy (up to 100 kHz) [11].

The human body generates alternating magnetic fields in the range of 0.1-0.2 mkT [12] simply by pumping blood. Since blood flow is facilitated through pulses, an alternating magnetic field can be produced around the blood vessels with a frequency 1-2 Hz. This is close to the values given in studies above. Thus, this “biogenic” alternating magnetic field can be harnessed and reapplied back to the body to produce the above-mentioned effects without the need for an external power source.

Therefore, the harnessed alternating magnetic field induced by the human blood flow can be reapplied back on to the blood vessels to improve blood circulation in the extremities.

To accomplish this the extremity can be encircled with wearable electroconductive coils. Because of the flow of blood an opposing alternating magnetic field (OAMF) will be induced around these coils as described by Lenz’s law. Just like the blood flow the OAMF will be modulated by repeating heart contractions (pulses). This will induce the OAMF in accord with the rate of blood flow, that is the blood flow will be influenced by the OAMF in the same modulation mode as delivered by blood flow pulse. This will have a direct effect on the blood flow and will expectedly filter out the influence of chaotic noise caused by external magnetic fields.

The largest contribution to OAMF flux density comes from the largest blood vessels going along the length of the extremities. In general, the direction of blood flow in veins is opposite to the direction of blood flow in arteries. Thus, with the flow of blood through the arteries an induced OAMF could assist the stagnating blood flow in veins to start moving faster. However, there are superficial veins in both upper and lower limbs that are close to the surface of the skin. The leg has small and great saphenous veins, whereas, the hand has cephalic, median cubital and basilic veins. Therefore, the magnetic field loss from these veins is the smallest.

The aim of the experiment was to find out if magnetic field generated by the human body affect the blood flow. For the first time was investigated the possible influence of metallized coils on human blood circulation. It was assumed that the current that is generated in the metallized coils is able to counteract the current that goes along with the bloodstream improving the blood circulation.

## Methods

### Rheovasography registration with metallized and non-metallized coils

Metallized coils were made of electroconductive material. Non-metallized coils were produced the same way of the same color as electroconductive coils to provide blind experiment and exclude placebo effect.

The experiment of possible influence of metallized coils on human blood circulation was divided into three parts: subject selection and data recording, experimental tests, signal processing.

For the experiment, the age group between 18 to 25 was chosen. This group consists of 12 students (7 male and 5 female) from Sports Academy of Latvia. Athletes have higher hemoglobin in compare to sedentary individuals. Due to this reason all tests were performed on athletes involved in endurance sports. All tests were made within the time of three month.

Experimental tests consist of two regimes – *physical movement* and *rest*. In physical movement regime subject was asked to move their leg with conductive coils on it up and down. In rest mode subject was asked to sit on the couch in comfortable position without moving their leg (see Table 1.).

**Table 1.**
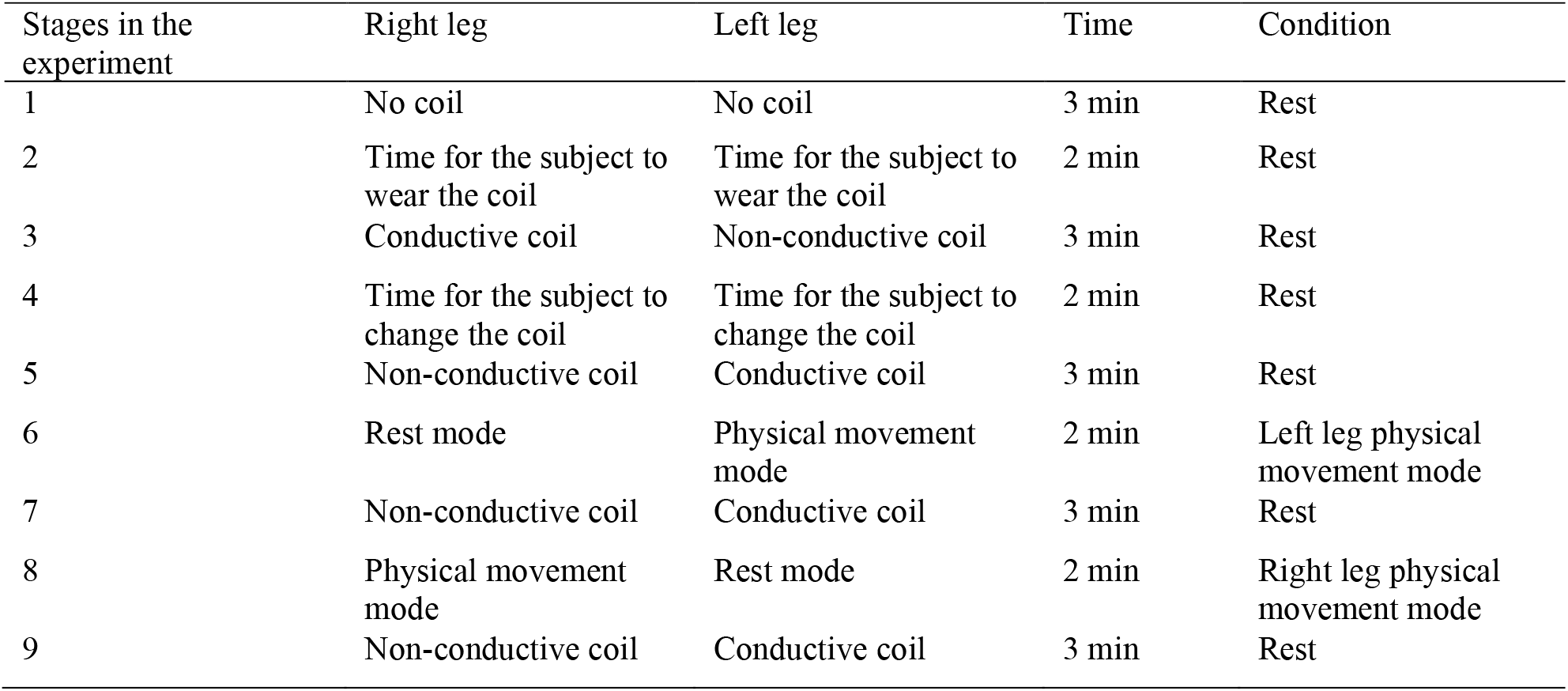
The algorithm of the experiment with non-conductive and conductive coils

| Stages in the experiment | Right leg | Left leg | Time | Condition |
| --- | --- | --- | --- | --- |
| 1 | No coil | No coil | 3 min | Rest |
| 2 | Time for the subject to wear the coil | Time for the subject to wear the coil | 2 min | Rest |
| 3 | Conductive coil | Non-conductive coil | 3 min | Rest |
| 4 | Time for the subject to change the coil | Time for the subject to change the coil | 2 min | Rest |
| 5 | Non-conductive coil | Conductive coil | 3 min | Rest |
| 6 | Rest mode | Physical movement mode | 2 min | Left leg physical movement mode |
| 7 | Non-conductive coil | Conductive coil | 3 min | Rest |
| 8 | Physical movement mode | Rest mode | 2 min | Right leg physical movement mode |
| 9 | Non-conductive coil | Conductive coil | 3 min | Rest |

Signal processing divides into *statistical analysis* and *data comparison*. For the statistical analysis recorded data was analyzed in different methods such as Gauss distribution, math expectation and histogram.

The Gaussian distribution shown is normalized so that the sum over all values of x gives a probability of 1. The nature of the Gaussian gives a probability of 0.683 of being within one standard deviation of the mean. The mean value is a=np where n is the number of events and p the probability of any integer value of x (this expression carries over from the binomial distribution). The standard deviation expression used is also that of the binomial distribution. [13]

To compare the data, a statistical assessment of changes in blood flow was performed. In the experiment mathematical expectation was used to determine Gaussian distribution. The goal was to overlap non-conductive and conductive signal so that the mathematical expectation is 0. If there is a difference in the signal, then mathematical expectation is not equal to 0. If there is a small difference in both negative and positive the resulting average will be close to zero. Metallized and non-metallized coils were compared in all combinations: left leg non-conductive and no coil; right leg non-conductive and no coil; left femur non-conductive and no coil; right femur non-conductive and no coil; left leg no coil and conductive coil; right leg no coil and conductive coil; left femur no coil and conductive coil; right femur no coil and conductive coil; left leg non-conductive and conductive coil; right leg non-conductive and conductive coil.

The rheovasogram analysis was conducted in *visual* and *digital analysis* steps. Visual analysis, using rheography-polyanalyzer made by Rean Poly, was based on interpretation of the external form of rheography wave, whereas digital analysis, using rheography-polyanalyzer, involves special digital calculations. In virtual analysis wave terminators should be defined at the beginning of the test (top and end). The area of the curve from the starting point to the top is called the ascending part of the anacrot, the area of the curve from the top to the end is called the descending or catacrota. [14] Digital analysis clarifies the nature of the visually observed changes and makes it possible to determine a number of features of the state of other vessels.

For rheovasography, four network devices are required: an electrocardiograph (ECG), special electrodes (strip electrodes), a rheography device and a computer for simultaneously displaying and recording a signal. This procedure uses a high frequency current that does not bring any unpleasant sensation to the subject. Blood can conduct an electric current, for this reason, the resistance varies in different phases of the blood flow. The dependence here is inversely proportional: the more the blood filling is, the less resistance of tissue appears. From high indicators to low there is a constant fluctuation. The rheovasogram graphically displays the passage of pulse, the increase of an amplitude when blood filling increases and, conversely, decrease of amplitude when blood filling decreases.

For the research, the patient lies on a couch, supine position. The skin of the extremities is degreased with alcohol. Sensors are attached to the treated areas. Signals are transmitted to the screen, the rheovasogram is recorded, the main indicators are calculated. The rheovasogram can be performed using multichannel rheography, or the investigation is carried out sequentially, starting from areas remote the center, ending with neighboring. A characteristic feature of the placement of conductors is strict symmetry. This method of data registration is non-invasive, and the patient is not subjected to any unpleasant or painful sensations, although rheovasography gives a reliable data. The patency of blood vessels, the quality of the outflow of blood, pathology and the level of their prevalence are determined with great accuracy. However, one of the methods disadvantages is a human factor. The ribbon electrodes must be placed in the same location for all the subjects. A small movement by the subject in measurement stage spikes a huge distortion which will take a few seconds to normalize. The quality of the result obtained can be affected by the κheographic attachments that can interfere.

To avoid placebo effect, in all tests similar non-conductive and conductive coil were used. It was not explained to the subjects about both coils and the idea of research before the experiment.

For rheovasography band electrodes are used, which are symmetrically applied to the left and right sides of the thigh and lower leg in the same areas of the skin. Low impedance in electrodes used for biopotential recordings in the experiment is required to ensure impedance matching with the environment and maximum energy transfer. This is to prevent reflections by the load - a bottleneck effect. The reason the kilo-Ohm is preferred for the input impedance is to keep the current in the circuit in the milli-Ampere range for electric safety. Blood has the least resistance, when an innocuous, weak current of high frequency (500 kHz and a small force of 10 μA) is passed through any section, then the electrical resistance of this section is recorded, which constantly changes. The curve of resistance changes reflects the blood filling of vessels when a pulse wave passes through them. Ribbon electrodes are called also band electrodes. The advantage of using a band electrode is, these are very efficient and can be easily fixed on the surface of the skin. Gel is used for binding the electrode to the subject’s skin and effectively dispersing electrical stimulation to the target area. [15] Ribbon electrodes were placed near the ankle, knee and femur areas throughout the experiment. The subjects had no itching sensation as gel was used to bind the electrode and skin. Even when changing coils, it was made sure that the electrode did not move and was placed in the same position.

The data acquisition algorithm was formulated for all subjects; in the same procedure; in the same set of combination with respect to time as shown in Table 1. Total time for one experiment on one subject is 23 minutes. After all the experiments the data was extracted from the rheovasograph and converted to Excel. Before the signal processing unwanted data was discarded; the stages when subject changes coils (stage 2 and 3) and physical movement stage (stage 6 and 8). Total data collected time is 15 min.

Electrodes were fastened below coils in both cases non-conductive and conductive (see Figure 1.).

**Fig. 1.**
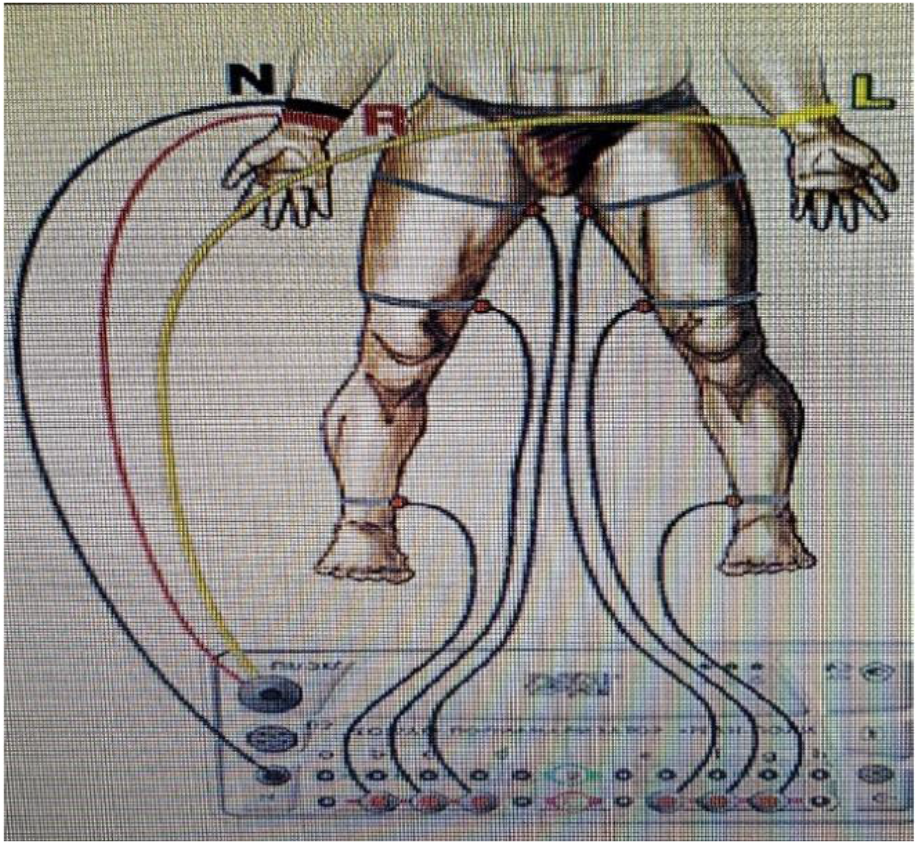
The electrode attachment places.

It was made sure that the band electrode is in constant contact with the skin. Before the experiment, subjects were asked not to make any movements during recording. The subject was placed in a dimly lit room on a comfortable couch, and the experimental area was adjusted to reduce the subject’s stress and anxiety.

## Results

### Rheovasogram analysis

To check the stability of the signal, covariance was applied. It was established that bioelectrical signal from vessels are weak and volatile. After checking the stability of the signal, following parameters were manually calculated: alpha (A) – duration of arterial inflow; beta (B) – duration of veins outflow; A1 (Ri) – rheographic index number of incoming blood (blood pulse and blood filling); A3 (Di) – diastolic index (outflow from the arteries to veins). As a signal is volatile six pulses were selected from each stage. It was decided to process the signal using different methods. For further signal processing different coil combinations were compared: no coil and non-conductive coil, no coil and conductive coil, non-conductive coil and conductive coil. Leg and femur were compared in different combinations graphically for all subjects. Angle and ratio for conductive and non-conductive cases were calculated, by calculating the width and height of the graphs.

Subtracted signal was processed with histogram distribution (see Figure 2). This process was carried for all combinations and all subjects. The amplitude differed when non-conductive and conductive were compared. To evaluate the difference between non-conductive and conductive coils the sharpness of diagram was calculated. To evaluate the sharpness of diagram maximum value (amplitude) and the width of histogram were taken.

**Fig. 2.**
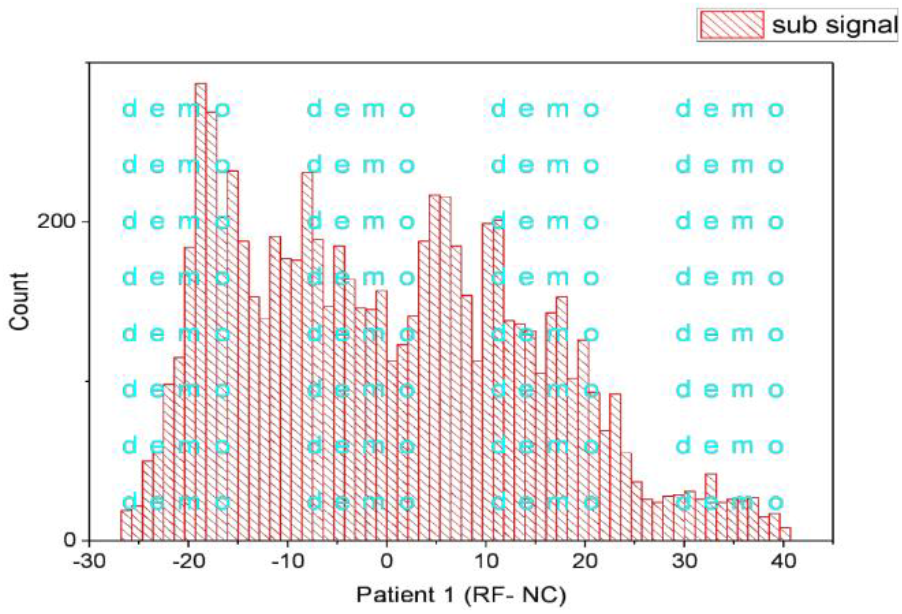
Non-conductive coil histogram for patient 1.

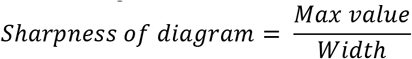

To determine the value of difference it was calculated.

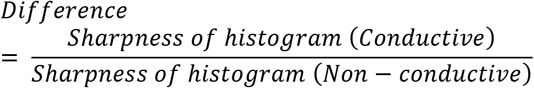

From all the histogram’s calculations, the difference of the sharpness of histogram is greater than 1 in all combinations (see Table 2.). This proves that there is a slight difference between conductive and non-conductive coil’s histogram. Conductive coil average maximum value is higher when compared to no-conductive coil. The sharpness of histogram is also higher, when load is applied. When non-conductive and non-conductive with load sharpness of diagrams are compared, the difference is noticeable. The same with conductive and conductive with load sharpness.

**Table 2.** The calculations of the sharpness of diagram achieved after rheovasography.

|  |  | Mean | SD |
| --- | --- | --- | --- |
| Right femur | Non-conductive | 8.65 | 6.87 |
|  | Conductive | 10.37 | 6.84 |
|  | Difference | 1.20 | 1.00 |
|  | Non-conductive | 6.12 | 3.47 |
|  | Non-conductive + load | 8.54 | 5.05 |
|  | Difference | 1.40 | 1.45 |
| Left femur | Non-conductive | 6.22 | 3.52 |
|  | Conductive | 8.53 | 5.32 |
|  | Difference | 1.37 | 1.51 |
|  | Conductive | 6.04 | 3.21 |
|  | Conductive + load | 8.47 | 3.67 |
|  | Difference | 1.40 | 1.15 |
| Right leg | Non-conductive | 2.20 | 0.68 |
|  | Conductive | 2.30 | 0.58 |
|  | Difference | 1.04 | 0.86 |
|  | Non-conductive | 2.55 | 0.83 |
|  | Non-conductive + load | 3.02 | 1.07 |
|  | Difference | 1.19 | 1.28 |
| Left leg | Non-conductive | 1.97 | 0.73 |
|  | Conductive | 2.32 | 0.85 |
|  | Difference | 1.17 | 1.17 |
|  | Conductive | 2.65 | 1.82 |
|  | Conductive + load | 3.37 | 2.00 |
|  | Difference | 1.27 | 1.10 |

Mathematical expectation was used to determine Gauss distribution. The objective was to overlap non-conductive and conductive signal so that the mathematical expectation is 0. If there is a difference in the signal, then mathematical expectation is not equal to 0. If there is a small difference in both negative and positive; the resulting average will be closer to zero. Both squared and absolute values was calculated for all subjects; for both femur and leg. Probability of significance of 95% for χ^2^ was considered. If χ^2^ > χ^2^_table value_, then the difference is significant. In all the combinations: non-conductive and conductive coils; no coil and conductive coil; non-conductive coil and no coil χ^2^ was greater than table value.

The OAMF induced by the flow of blood through the larger veins and arteries has a non-zero value and thus could affect the flow of blood in veins. A simplified vascular model of the lower leg was made. The total magnetic induction (B) caused by blood flow and situated around the vessels is a product of superposition of induction produced by blood flow in arteries (B_a_) and veins (B_v_):

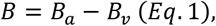

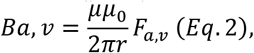

a, v – indexes related to the flows in arteries and veins, respectively; *μ, μ*_0_ – magnetic permeability of the leg tissue and magnetic constant, respectively; F_a,v_ – total blood flow in the leg produced by both arteries and veins; r – the distance from the blood vessel to the coil located around the leg.

The lower leg has three main arteries. The posterior tibial artery and the tibial artery are located at around ¼ of the leg radius from the center of the lower leg while the anterior tibial artery is located at about ½ the leg’s radius from the center of the leg.

The lower leg also has eight main veins. The two small saphenous veins and both the anterior and the posterior accessory great saphenous vein are located close to the surface of the leg at about one lower leg radius from the center of the leg. The posterior tibial vein and the fibular vein are located close to the center of the lower leg at about ¼ of the lower leg’s radius. The anterior accessory great saphenous vein and the great saphenous vein are located at about ½ the leg’s radius.

All the arteries and veins proceed along the length of the tibia and terminate at the lower end of the bones. Both the main arteries and the main veins are situated somewhat parallel to each other going along the leg.

When blood is being pumped through the arteries and veins the sum magnetic induction (B) caused by their flow and situated around the vessels is the product of superposition of the induced fields produced by blood flow in arteries (B_a_) and veins (B_v_). Therefore, an approximate value for B in the center of the leg could be written down as follows:

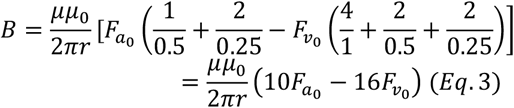

F_a0_, F_v0_ – blood flow in a single artery and vein, respectively. As indicated in [16] F_a0_ and F_v0_ are ∼ 16.5 and ∼0.2 cm3/sec, respectively. Putting these values into Eq. 3:

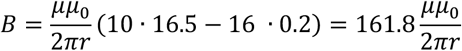

Though Eq. 3 is a rough estimate, it shows that B≠0, thus induction can occur. Of course, more complex models need to be developed for a more precise description of the above-discussed EM phenomena. However, this still provides credence to the idea of employing the magnetic field produced by the blood flow in the leg for OAMF induction in an external coil.

## Discussion

To the best of our knowledge, our study is the first to investigate how the metallized coils can affect the human blood circulation. This happens because of the OAMF induced in the coils, as described by Lenz’s law.

These findings were surprising, since the human body generated alternating magnetic fields are in the range of 0.1-0.2 mkT [12] and for the magnetotherapy procedure complicated and massive devices are required. [11] However, it is assumed that from the human body induced OAMF in metallized coils can have a prolonged effect on the blood circulation. Metallized coils are easy and comfortable to wear. Therefore, conductive coils can be used as a gentle treatment or for rehabilitation purposes.

The study by Gutiérrez-Mercado et al [6] has similar idea by using low magnetic fields for treatment; however, this study was conducted on rats and no possible low magnetic field influence on human blood circulation was investigated. In this study an experimental group subjected to extremely low frequency magnetic fields (120 Hz harmonic waves and 0.66 mT) by the use of Helmholtz coils.

Another study that shows the positive influence of pulsed electromagnetic fields on blood circulation was conducted by Lee et al [7]. This study on rats shows that the PEMF (1 Hz, 10 mT) may improve blood circulation. However, in our study was shown that the by the human body generated alternating magnetic field in the range of 0.1-0.2 mkT can influence the blood circulation. This assumption is underpinned by the difference in the diagram sharpness combining non-conductive and conductive coils. The sharpness of the diagram between right femur non-conductive (8.65±6.87) and conductive (10.37±6.84) coils, and left femur non-conductive (6.22±3.52) and conductive (8.53±5.32) differs. This shows that there is a difference in blood circulation, when conductive coils are on.

## Conclusion

The aim of the experiment was to find out if metallized coils affect the blood flow. For the first time was investigated the possible influence of metallized coils on human blood circulation. It was assumed that the current that is generated in the metallized coils is able to counteract the current that goes along with the bloodstream improving the blood circulation.

It has been achieved that metallized coils affect the blood flow, because of the difference between the sharpness of the diagrams. Difference between the sharpness of the diagram: for right femur between non-conductive and conductive coil is 1.20±1.00; for left femur between non-conductive and conductive coil is 1.37±1.51; for right leg between non-conductive and conductive coil is 1.04±0.86; for left leg between non-conductive and conductive coil is 1.17±1.17. However, the difference between the sharpness of the diagram: for right femur between non-conductive and non-conductive with load coil is 1.40±1.45; for left femur between conductive and conductive with load coil is 1.40±1.15; for right leg between non-conductive and non-conductive with load coil is 1.19±1.28; for left leg between conductive and conductive with load coil is 1.27±1.10. If compare the differences in the sharpness of diagrams conductive coils provide effect as if leg was loaded; this claims that metallized coils has influence on human blood circulation. Furthermore, Gaussian distribution points out that there is an increase in all cases of data when compared to the table value; this shows that there is a difference in signal as it is not equal to zero.

We suggest that metallized coils could help to treat patients with CVI and varicose veins. Our results may serve to stimulate future investigations into the possible effects of metallized coils on different age groups. This could be considered as a gentle training of the human circulatory system.

## Abbreviations

RI: Rheography index
DI: Diastolic index
OAMF: Opposing alternating magnetic field
CVI: Chronic venous insufficiency
ECG: Electrocardiography
PEMF: Pulsed electromagnetic field
CY: Cycle

## References

1. Young JY, Juyong L (2018) Chronic venous insufficiency and varicose veins of the lower extremities 34(2): 269–283 https://dx.doi.org/10.3904%2Fkjim.2018.23 0

2. Eberhardt RT., (2014) Raffetto JD., Chronic Venous Insufficiency, Circulation 130:333–346. https://doi.org/10.1161/CIRCULATIONAH A.113.006898

3. Rabe E, Guex JJ, Puskas A, et al (2012) Epidemiology of Chronic Venous Disorders in Geographically Diverse Populations: Results From the Vein Consult Program, Int Angiol 31(2):105–15.

4. Gasper W.J., Grenon M., Hiramoto J.S., et al Chronic Venous Insufficiency, University of California San Francisco, Department of Surgery https://surgery.ucsf.edu/conditions--procedures/chronic-venous-insufficiency.aspx

5. Szopinski JZ (2014) The Biological Action of Physical Medicine: Controlling the Human Body’s Information System 244.

6. Gutiérrez-Mercado YK, Cañedo-Dorantes L, Gómez-Pinedo U et al (2012) Increased Vascular Permeability in the Circumventricular Organs of Adult Rat Brain Due to Stimulation by Extremely Low Frequency Magnetic Fields, Bioelectromag 34(2):145–55 https://doi.org/10.1002/bem.21757

7. Lee N, Park J, Choi Y et al (2018) Effect of pulsed electromagnetic fields on the blood circulation in ischemic skin models: A pilot study J Electromag Biol Med 37:202–207. https://doi.org/10.1080/15368378.2018.1523798

8. McNamee DA, Legros AG, Krewski DR et al (2009) A Literature Review: The Cardiovascular Effects of Exposure to Extremely Low Frequency Electromagnetic Fields Int Arch Occup Environ Health 82(8):919–33 https://doi.org/10.1007/s00420-009-0404-y

9. Nishimura T, Tada H, Guo X et al (2011) A 1-μT Extremely Low-Frequency Electromagnetic Field vs. Sham Control for Mild-To-Moderate Hypertension: A Double-Blind, Randomized Study, Hypertens Res 34(3):372–7 https://doi.org/10.1038/hr.2010.246

10. Peter Bednarčík (2019) Principle of Pulse Electromagnetic Field Therapy (PEMF) https://www.biomag-medical.com/about-magnetotherapy/

11. Peter Bednarčík (2019) Devices and procedures using an EMG field and electric energy https://www.biomag-medical.com/about-magnetotherapy/devices-and-procedures-using-an-emg-field-and-electric-energy/

12. Gross S, Barmet C, Dietrich BE et al (2016) Dynamic nuclear magnetic resonance field sensing with part-per-trillion resolution, Nature Communications 7:13702 https://doi.org/10.1038/ncomms13702

13. Gaussian distribution function http://hyperphysics.phy-astr.gsu.edu/hbase/Math/gaufcn.html

14. The Institute of Traumatology and Orthopedics Reovasography https://ito.gov.ua/en/our-services/diagnostics/reovasography.html

15. Klett J (2014) Why Electrode Gel is Key to Successful Electrotherapy http://www.therasigma.com/why-electrode-gel-is-key-to-successful-electrotherapy/

16. Anatomy and Physiology II, The Cardiovascular System: Blood Vessels and Circulation, Blood Flow, Blood Pressure, and Resistance https://courses.lumenlearning.com/ap2/chapter/blood-flow-blood-pressure-and-resistance-no-content/

17. Ivanov AI, Kawamoto H. Sankai Y(2012) Development of a capacitive coupling electrode for bioelectrical signal measurements and assistive device use https://doi.org/10.1109/ICCME.2012.62757 20

18. Van den Oever R, Hepp B, Debbaut B, et al (1998) Socio-economic Impact of Chronic Venous Insufficiency. An Underestimated Public Health Problem, Int Angiol. 17(3):161–7.

19. Beebe-Dimmer JL, Pfeifer JR, Engle JS (2005) The Epidemiology of Chronic Venous Insufficiency and Varicose Viens, Ann Epidemiol 15(3):175–84. https://doi.org/10.1016/j.annepidem.2004.05.015

20. Carpentier PH, Maricq HR, Biro C et al (2004) Prevalence, risk factors, and clinical patterns of chronic venous disorders of lower limbs: A population-based study in France, J Vasc Surg 40:650–659 https://doi.org/10.1016/j.jvs.2004.07.025

21. Chiesa R, Marone EM, Limoni C et al (2005) Chronic Venous Insufficiency in Italy: The 24-cities Cohort Study, Eur J Vasc Endvasc Surg 30:422–429. https://doi.org/10.1016/j.ejvs.2005.06.005

22. Ross CL, Ang DC, Almeida-Porada G (2019) Targeting Mesenchymal Stromal Cells/Pericytes (MSCs) With Pulsed Electromagnetic Field (PEMF) Has the Potential to Treat Rheumatoid Arthritis, Front Immunol 10:266 https://doi.org/10.3389/fimmu.2019.00266

23. Chung YH, Lee YJ, Lee HS et al (2014) Extremely Low Frequency Magnetic Field Modulates the Level of Neurotransmitters, Korean J Physiol Pharmacol 19(1):15–20 https://dx.doi.org/10.4196%2Fkjpp.2015.19.1.15

